# A novel druggable DYRK3/CAMKV signaling module for Neuroblastoma tumor growth inhibition

**DOI:** 10.1101/2023.11.30.569438

**Authors:** Esteban J. Rozen, Kim Wgglesworth, Jason Shohet

## Abstract

High-risk Neuroblastoma is a very aggressive and deadly pediatric cancer, accounting for over 15% of all childhood cancer mortality. Therefore, novel therapeutic strategies for the treatment of neuroblastoma are urgently sought for. Here, we identified the DYRK3 kinase as a critical mediator of neuroblastoma cell proliferation and *in vivo* tumor growth. Our data suggest a role for DYRK3 as a regulator of the neuroblastoma-specific protein CAMKV, which is also required for neuroblastoma cell proliferation. We show that CAMKV is phosphorylated by DYRK3, and that inhibition of DYRK3 kinase activity induces CAMKV aggregation, probably mediated by its highly disordered C-terminal half. Importantly, we provide evidence that the DYRK3/CAMKV signaling module could play an important role in the regulation of the mitotic spindle during cell division, supporting the idea that inhibition of DYRK3 and/or CAMKV in neuroblastoma cells could constitute an innovative and highly specific intervention to fight against this dreadful pediatric cancer.

## Introduction

Neuroblastoma (NB) is the most common extracranial solid tumor of the childhood, causing about 15% of all pediatric cancer deaths^1^. Despite very aggressive multimodal interventions, only ∼50% of high-risk NB patients survive, exhibiting serious long-term sequelae from therapy. Therefore, more efficient, and less toxic interventions are urgently needed. In search of potential druggable targets in NB pathogenesis, we performed a systematic analysis of the correlation between expression of the Dual-specificity tYrosine phosphorylation-Regulated Kinase (DYRK) family members and neuroblastoma (NB) patient survival probability. We annotated and ranked the expression of all 5 DYRK kinases (DYRK-1A, 1B, 2, 3 and 4) across 10 publicly available NB patient datasets from the ‘R2: Genomics Analysis and Visualization Platform’ (as previously described^2^) and observed a significant and robust correlation between DYRK3 expression and a worse patient survival in 8 of these 10 cohorts, but not for any other family members (Table 1). This observation was also true for other neurological tumors (particularly Glioma, supplementary fig. 1).

**Table 1.**
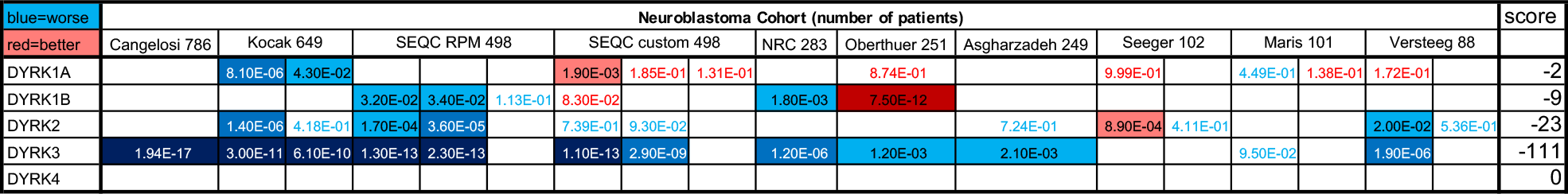
Bonferroni adjusted p-values for the correlations between DYRK family members expression and NB patient survival probability. Values highlighted in blue denote significant negative correlations (higher expression = reduced survival probability ‘worse’), while values highlighted in red denote positive correlations (higher expression = increased survival probability ‘better’). Blue/red fonts denote non-significant values (bonf. p-val > 5.00E-02). Scores are arbitrary units based on the number of datasets showing one or more significant correlations for a given gene, and the value of such significance. We arbitrarily define that a gene must be significantly correlated to either a better or a worse outcome in at least 5 independent cohorts, and obtain a minimum of 50 points (either positive or negative, respectively), to be considered a significant hit.

Increased DYRK3 expression has been associated with a higher aggressiveness of Glioblastoma cells and tumors^3^, although another report suggested it might negatively regulate hepatocellular carcinoma progression^4^. Noteworthy, Ivanova *et al*.^5^ reported a role for DYRK3 as a negative regulator of the hypoxic response and differentiation in NB cells, hence contributing to their aggressive behavior. DYRK3 remains largely understudied, although recent reports from Lucas Pelkmans’ group and others have implicated this kinase as a critical regulator of several **biomolecular condensates** (BCs), such as Stress Granules^6,7^, centrosomes and the mitotic spindle^8^, endoplasmic reticulum exit sites^9^ or Mediator complex condensates^10^. BCs -also referred to as membranelles organelles- are sub-cellular compartments that form through liquid-liquid phase separation of RNA and/or proteins that contain intrinsically disordered regions (IDRs)^11,12^. Such IDRs can undergo numerous post-translational modifications, particularly phosphorylation, which regulate their partitioning into these condensates^13–15^.

## Results

To investigate in more detail the relation between DYRK3 levels and neuroblastoma pathophysiology, we knocked-down its expression by means of 3 specific shRNA-expressing lentiviral clones (shDYRK3-1/2/3) vs. a non-targeting control (shCtrl) in 2 different NB cell lines. In this way, we observed a very striking impairment in NB cell proliferation (Figure 1A). Efficient and specific down-regulation of DYRK3 was confirmed by quantitative Real Time PCR (RT-qPCR; Figure 1B and supplementary fig. 2), demonstrating a critical role for DYRK3 in neuroblastoma cell homeostasis. To further examine DYRK3 implications in NB tumorigenesis, we established a tetracycline-inducible shDYRK3 expression system (Tet-shDYRK3 JF cells). Validation of this paradigm by RT-qPCR and Western Blot (WB) confirmed efficient downregulation of DYRK3 upon doxycycline treatment (Figure 1C and supplementary fig. 3), again resulting in significantly impaired cell proliferation (Figure 1D). Treatment of JF NB cells with low doses (1 µM) of Harmine, a pan-DYRK inhibitor known to block DYRK3, resulted in a moderate but significant inhibition of cell proliferation (supplementary fig. 4). At higher doses (10 µM) all cells died within 48 hours (not shown), suggesting a non-specific toxic effect.

**Figure 1.**
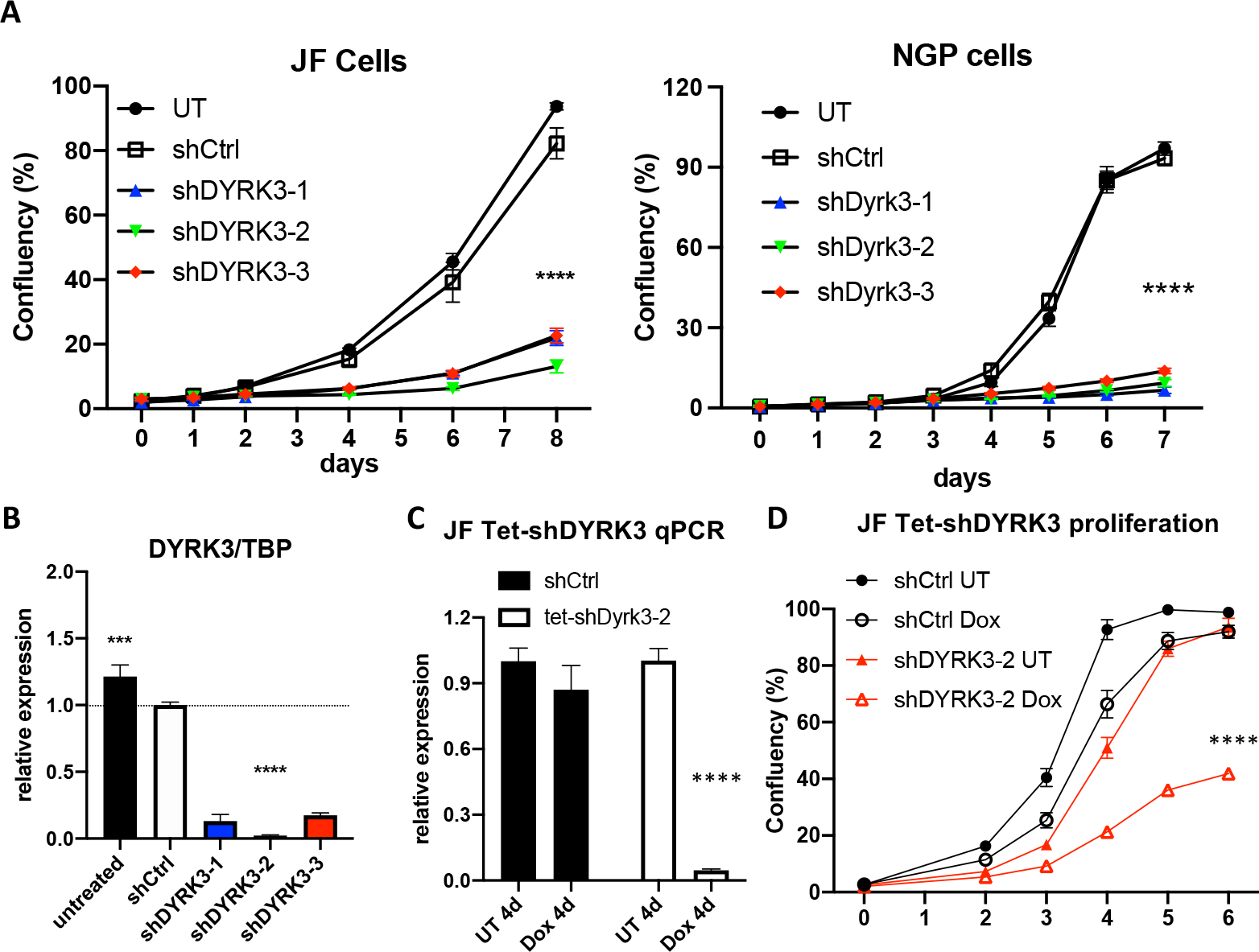
**A**. NGP and JF neuroblastoma cells were left untreated (UT) or transduced with lentiviral vectors for expression of the indicated shRNAs. After puromycin selection, ∼2×10^3^ cells were plated on M96 well plates (4 replicates/condition) and allowed to grow for 8 days. Confluency of each well was measured in an Image cytometer (Celigo, Nexcelom). **B**. JF cells form A were collected in triplicate and processed for RNA extraction and qRT-PCR analysis of DYRK3 expression (vs. the housekeeping gene TBP). **C**. Tetracycline-inducible shRNA system validation by qRT-PCR for DYRK3/TBP expression demonstrated efficient and specific DYRK down-regulation upon doxycycline treatment of Tet-shDYRK3-expressing cells. **D**. Tet-shRNA-expressing cells (shCtrl vs. shDYRK3) were plated on M96 well plates (4 replicates/ condition) and allowed to grow for 6 days in absence or presence of doxycycline. Confluency of each well was measured as in A.

To assess the direct role of DYRK3 in NB tumorigenesis *in vivo*, Tet-shDYRK3 JF cells were orthotopically injected into the renal capsule of immuno-compromised NSG™ recipient mice. Seven days post-surgery (7 dps), mice were randomly divided into ‘control’ (untreated) group vs. ‘doxy’ group, receiving 1mg/ml doxycycline in the drinking water for 3 additional weeks (Figure 2A). Importantly, all mice in the control group developed large tumors, while the doxy group tumors were dramatically smaller (Figure 2B-C). As a control, the contralateral healthy kidneys showed no significant weight differences (supplementary fig. 5). RT-qPCR analysis from tumor RNA samples reflected the efficient downregulation of DYRK3 by doxycycline treatment (Figure 2D), thus confirming a critical and previously unrecognized role for this kinase in NB cell proliferation and *in vivo* tumor growth. As mentioned above, Ivanova *et al*.^5^ reported a role for DYRK3 as a negative regulator of the hypoxic response and differentiation in NB cells, hence contributing to their aggressive behavior. Surprisingly, this work was entirely based on DYRK3 endogenous or ectopic expression, but no pharmacological inhibition or downregulation was performed, and did not examine potential effects on NB cell proliferation. Hence, our robust preliminary results are not in disagreement with such a function in a hypoxic setting.

**Figure 2.**
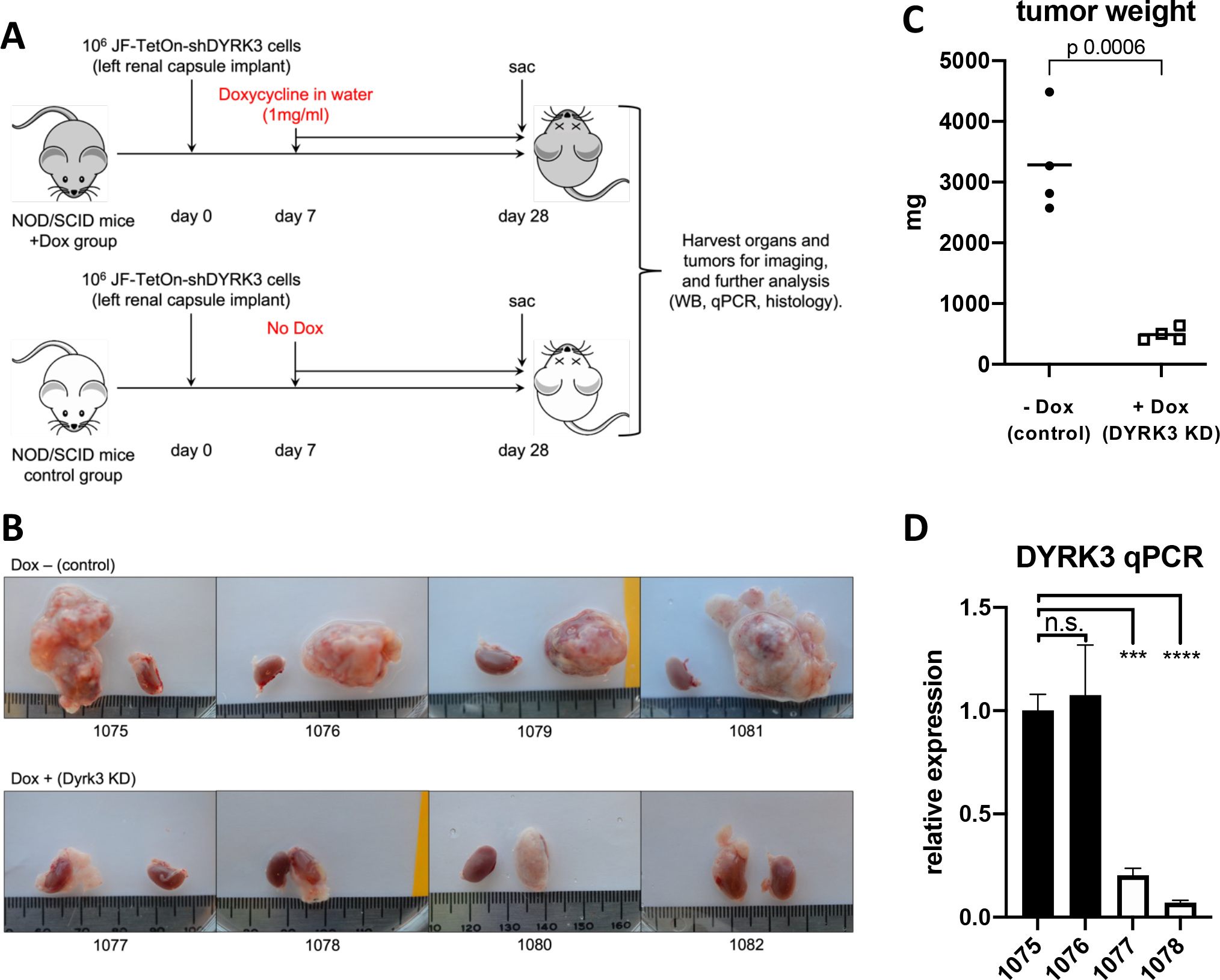
**A**. Experimental design for assessment for *in vivo* examination of DYRK3 implications on NB tumor growth. **B**. Pictures of JF-derived renal tumors and corresponding contralateral kidneys (as control) at experimental endpoint (28 dps), showing the striking defect on tumor growth by Tetracycline-inducible DYRK3 downregulation (n= 4 animals/group; see main text). **C**. Tumor growth inhibition was quantified as tumor weight, showing a very significant reduction in tumor mass by DYRK3 downregulation **D**. RNA was isolated from 2 tumors per group and subjected to qRT-PCR to quantify DYRK3 expression, demonstrating efficient DYRK3 downregulation by doxycycline treatment.

In search of a mechanistic target, we explored the literature for known DYRK3 substrates that could have a specific role in NB tumorigenesis. Wippich *et al*.^6^ performed an *in vitro* kinase substrate identification screen using protein microarrays in the presence of wild-type (WT) recombinant DYRK3 vs. a kinase dead mutant (K128M), as a negative control. Among the 26 protein target hits phosphorylated only by WT DYRK3, they found FIP1L1, AKTS1S and CAMKV as the top 3 candidates by average Z-score. The authors went on to characterize AKT1S1 (PRAS40) as a novel phosphorylation substrate of DYRK3 in stress granule biology. We became interested in CAMKV, a protein that is highly enriched in neuroblastoma cancer cell lines (Figure 3A) and in healthy neural tissues (Figure 3B). Importantly, CAMKV expression has previously been correlated to a worse NB patient survival and its expression is highly associated to MYCN or MYC expression in NB cell lines and primary tumor samples^16^. Therefore, we decided to explore in more detail the potential relation between DYRK3 and CAMKV in NB cell homeostasis. To corroborate that CAMKV indeed can interact with DYRK3, we used NGP NB cells transduced with a lentiviral vector to stably over-express an mCherry-CAMKV fusion protein. These cells were transiently transduced with an HA-tagged DYRK3 WT (HA-DYRK3) construct or an empty vector as control, and the corresponding cell lysates were subjected to immuno-precipitation (IP) with an mCherry-specific antibody. Western blot analysis of these immuno-precipitates revealed a clear anti-HA signal in the HA-DYRK3-tranfected sample, but not in the control, demonstrating DYRK3 coprecipitation with CAMKV (Figure 3C). Finally, mCherry immuno-precipitates from NGP mCherry-CAMKV cells were subjected to *in vitro* kinase assays (IVKs) with radioactive [γ-^32^P]-ATP, in the absence or presence of recombinant human DYRK3. This approach confirmed the direct *in vitro* phosphorylation of CAMKV by DYRK3 (Figure 3D).

**Figure 3.**
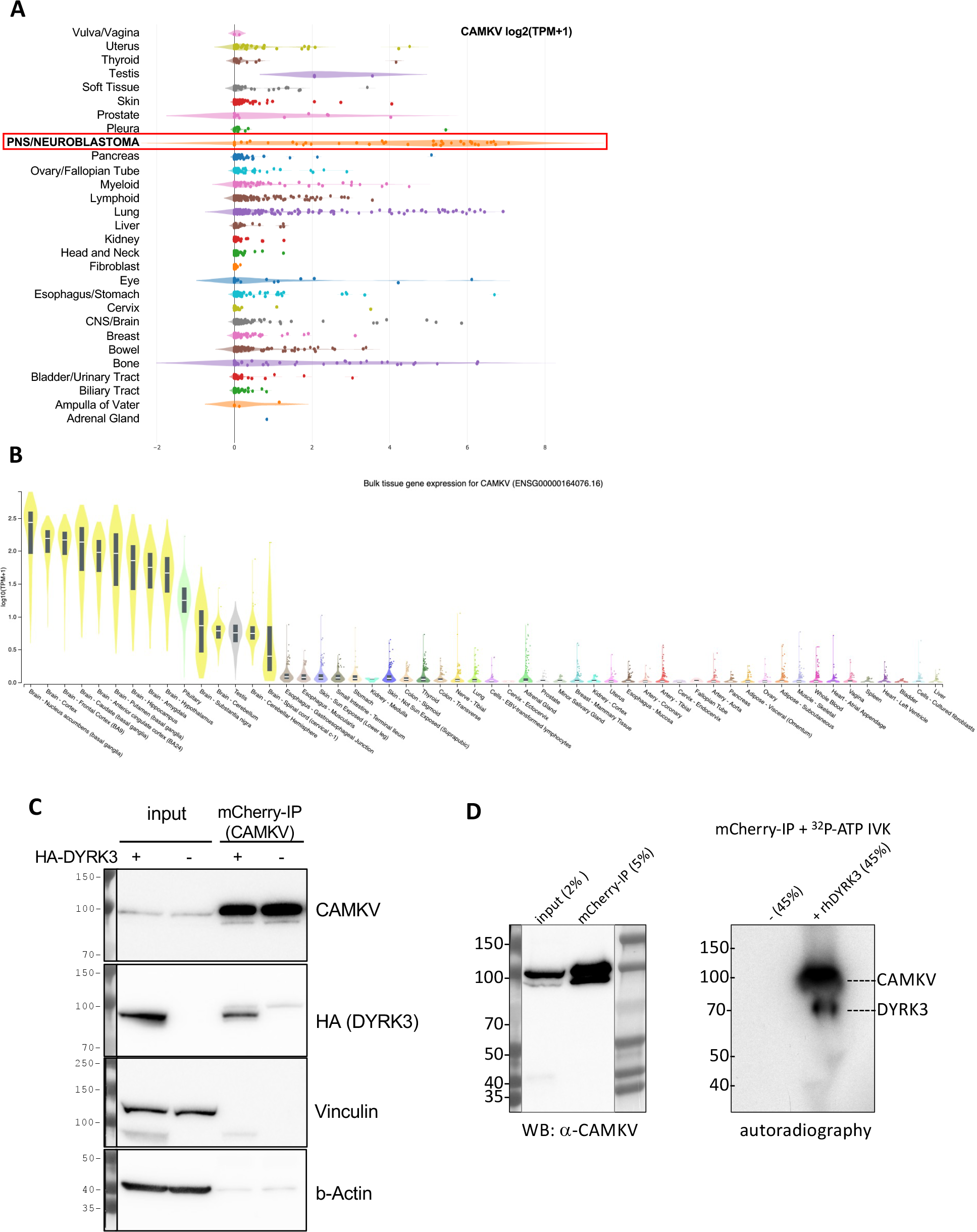
**A**. Expression of CAMKV across different cancer cell lines from the depmap portal (depmap.org) demonstrates a high and specific expression in neuroblastoma cell lines as compared to most other cancer cell lines. **B**. Analysis of CAMKV expression across human tissues from the Genotype-Tissue Expression Portal (gtexportal.org) demonstrates that CAMKV expression is exclusively enriched in brain tissues. **C**. Immunoprecipitation studies in NGP NB cells overexpressing mCherry-CAMKV WT +/- HA-DYRK3 demonstrate DYRK3 (HA) co-precipitation with mCherry (CAMKV) immuno-complexes. **D**. *In vitro* kinase assay with radioactive [γ-^32^P]-ATP, in the absence or presence of recombinant human DYRK3 confirmed the direct *in vitro* phosphorylation of CAMKV by DYRK3.

Given the known roles for DYRK3 as a critical regulator of the liquid-liquid phase separation (LLPS) behavior of its substrates, we then focused on CAMKV’s protein sequence. Barylko *et al*.^17^ recently reported a predicted intrinsically disordered region (IDR) of ∼200 amino-acids in CAMKV’s C-terminal half. We employed bioinformatic tools for the prediction of LLPS formation - ParSe (phase-separating protein regions prediction tool; Figure 4A)- or for prediction of disordered regions -PONDR^®^ (Predictor of Natural Disordered Regions) and PrDOS (Protein DisOrder prediction System; supplementary fig. 6)-, which further confirmed the presence of a highly disordered C-terminal region likely capable of undergoing phase separation in a phosphorylation-regulated manner. Upon a deeper look, we identified 7 tandem repeats of an octapeptide motif (D-X-X-X-T-P-A-T), including 2 canonical and 5 highly related DYRK phosphorylation motifs (Figure 4B). In fact, in the original characterization of rat Camkv, Godbout *et al*.^18^ briefly described such region as a potential ‘PEST sequence’, a motif known to act as a signal for degradation in proteins with a short half-life^19,20^. In this context, our overexpression experiments with a lentivirally encoded mCherry-CAMKV WT construct (see below) suggested CAMKV as a very stable protein, as demonstrated by the high and constant expression of mCherry-CAMKV across many passages with no loss (but rather increase) of the signal along time (not shown). Furthermore, in an attempt to characterize the role of this unique sequence, we generated an overexpression mutant version, ‘mCherry-CAMKV-ΔIDR’, lacking the 56 amino-acids corresponding to the 7 tandem octapeptide repeats. Surprisingly, the mutant variant exhibited a very low to null expression, suggesting that the resulting protein was very unstable (Figure 4C), and confirming that the 7 tandem octapeptide repeats are needed for expression or stabilization -and likely function- of CAMKV, and not acting as a canonical PEST sequence. Interestingly, shRNA-mediated downregulation of CAMKV by shRNA lentiviral vectors also resulted in a striking impairment of NB cell proliferation (Figure 4D). Of note, the level of CAMKV downregulation by different shRNA efficiencies (Figure 4E) was nicely correlated to the degree of proliferation potential of these cells (Figure 4D), suggesting that CAMKV is required for NB cell proliferation or survival in an expression level-dependent fashion.

**Figure 4.**
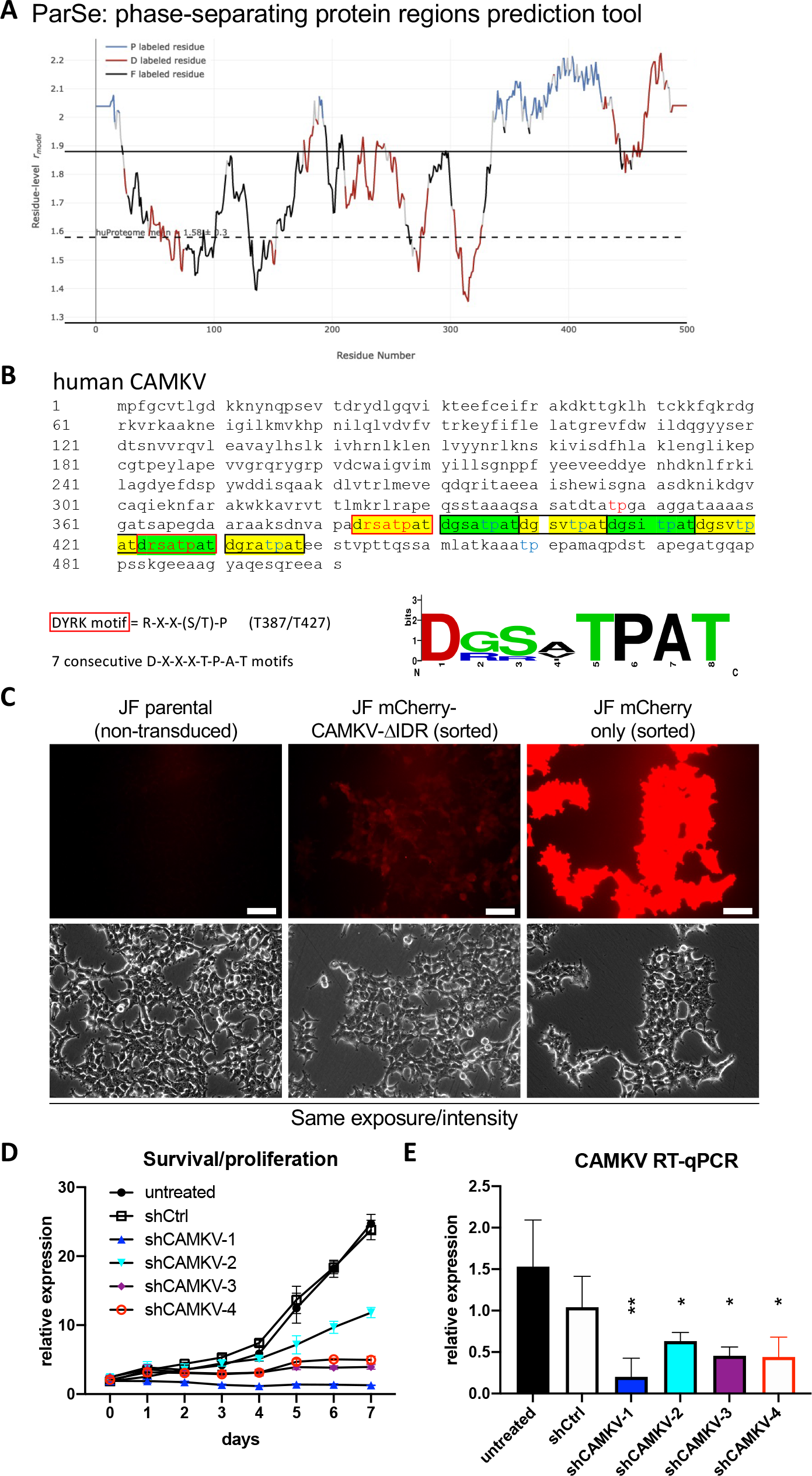
**A**. CAMKV protein sequence was analyzed with the phase-separating protein regions online prediction tool, ParSe. P Regions (P) are intrinsically disordered and prone to undergo LLPS. D Regions (D) are intrinsically disordered but do not undergo phase separation. F Regions (F) may or may not be intrinsically disordered, but can fold to a stable conformation. **B**. CAMKV protein sequence depicting the 7 tandem octapeptide motifs (D-X-X-X-T-P-A-T), including 2 canonical DYRK phosphorylation sites (T387 and T427, red fonts). **C**. JF neuroblastoma cells were left untreated (parental) or transduced with lentiviral vectors for overexpression of the indicated construct. Cells were then sorted for purity. Cells tranduced with mCherry-CAMKV-ΔIDR had a very low mCherry signal, suggestive of a highly unstable product. Scale bar = 40 µm **D**. JF cells were left untreated or transduced with indicated shRNA-expressing lentiviral vectors. Cells were selected with puromycin for 3-4 days and then subjected to time-course proliferation assays (4 replicates/ condition). Confluency of each well was measured in an Image cytometer (Celigo, Nexcelom), demonstrating a severe impairment in cell proliferation by CAMKV downregulation. **E**. The same cells used in D were processed in parallel for RNA extraction and subjected to RT-qPCR to confirm CAMKV mRNA downregulation.

To examine whether CAMKV might form or localize to membraneless organelles, we analyzed our mCherry-CAMKV overexpressing cells. CAMKV was originally predicted to lack a kinase activity and associated with neuronal vesicles of the rat cortex^18^. More recent work in mouse models implies a role in activity-dependent bulk endocytosis during the recycling of synaptic vesicles^21^, while Liang *et al*.^22^ suggested the co-localization of Camkv with postsynaptic scaffold protein PSD-95 puncta, where it may be required for the activity-dependent maintenance of dendritic spines. Nevertheless, in those immunofluorescent staining images, Camkv was not specifically localized to the dendritic spines, but rather homogeneously distributed all along the neuron. Additionally, Barylko *et al*.^17^ found that murine Camkv can undergo palmitoylation on its N-terminal end and that this modification was necessary for plasma membrane localization of a Camkv-EGFP (C-terminal) fusion construct. Interestingly, the authors noted that an N-teminal fusion construct (EGFP-Camkv) did not localize to the membrane, but had a homogeneous cytosolic distribution. As for human CAMKV, Sussman *et al*.^16^ also suggested a membrane localization in NB cell lines, although their data was largely inconclusive. In our hands, ectopic expression of mCherry-CAMKV in NB cell lines showed a clear homogeneous cytosolic pattern (Figure 5A), consistent with that reported by Barylko *et al*.^17^ for EGFP-Camkv. More importantly, immuno-fluorescent staining of endogenous CAMKV confirmed a similar cytoplasmic distribution in interphasic NB cells (Figure 5B). Since inhibition of DYRK3 has been shown to affect the organization of several biomolecular condensates^6–10^, we treated mCherry-CAMKV NGP cells with Harmine, a pan-DYRK inhibitor known to block DYRK3. Interestingly, this treatment resulted in the relocalization of mCherry-CAMKV into numerous aggregates (Figure 5C and supplementary fig. 7) with a very dynamic behavior (supplementary videos 1 and 2). Treatment of these cells with an unrelated DYRK3 inhibitor, GSK-626616, showed similar results (supplementary fig. 8). These results are consistent with previous observations in other biomolecular condensates upon DYRK3 inhibition^6,8,9^, and suggest that CAMKV might indeed undergo liquid-liquid phase separation in a DYRK3-regulated fashion. Noteworthy, we failed to observe aggregation of endogenous CAMKV upon Harmine treatment (not shown). This might suggest that the mCherry-CAMKV aggregates are a result, at least in part, from its non-physiological over-expression, or that endogenous CAMKV aggregates are sensitive to the harsh methanol fixation used for this staining. Importantly, in our immuno-fluorescence staining analyses of endogenous CAMKV we noticed that virtually every NB cell undergoing cell division displayed a considerably higher anti-CAMKV signal. Surprisingly, this signal corresponded to a very clear staining of the mitotic spindle (Figure 5D and supplementary fig. 9), a transitory structure fundamental for the progression of the cell cycle whose organization is governed by liquid-liquid phase separation^23–25^. As mentioned above, Rai *et al*.^8^ reported DYRK3 colocalization with Pericentrin in the mitotic spindle poles, further supporting a potential direct role in the regulation of CAMKV function during cell division. Interestingly, we also observed a clear mitotic spindle pole localization of endogenous phospho-AKT1S1 (Thr246) in dividing JF cells (Figure 5E left panel). As mentioned above, Wippich *et al*.^6^ characterized the direct phosphorylation of Thr246-AKT1S1 by DYRK3, although a function for this signaling module in the regulation of the mitotic spindle has never been reported. Importantly, we further noticed that AKT1S1 expression is also highly correlated to a worse NB patient outcome (Figure 5E right panel), as it has been suggested in several other cancer types^26^. This observation supports the idea that DYRK3 might act as a central orchestrator of the mitotic spindle organization by recruiting and/or phosphorylating CAMKV and AKT1S1. Therefore, we hypothesize that down-regulation or pharmacologic inhibition of DYRK3, CAMKV and AKT1S1 in NB cells results in an impairment of the mitotic spindle organization and subsequent exit from the cell cycle, a function that will be the focus of future efforts.

**Figure 5.**
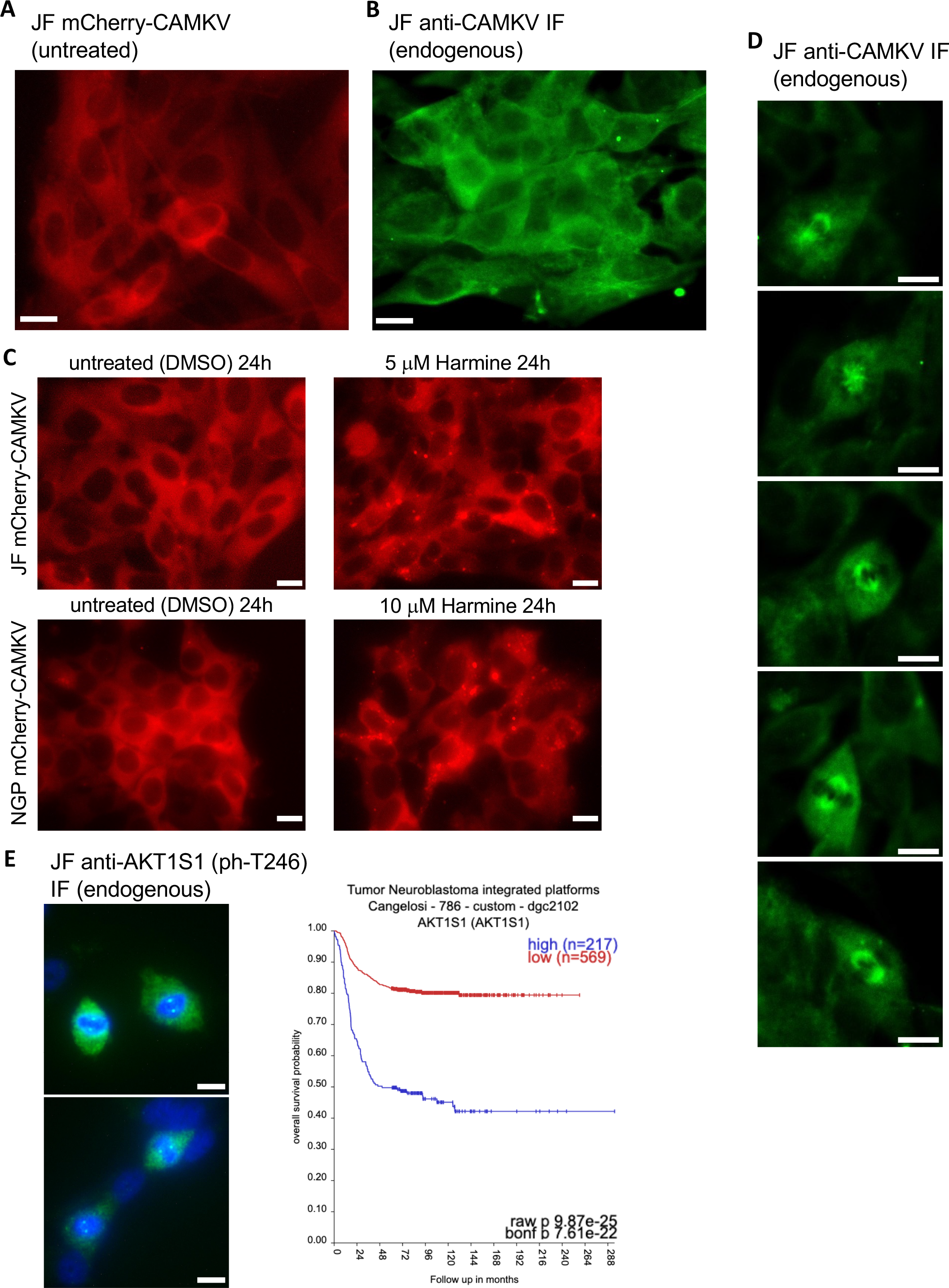
**A**. Fluorescence image of live JF cells transduced with a lentiviral vector for expression of mCherry-CAMKV (WT), demonstrating a homogeneous cytosolic distribution of the mCherry signal. **B**. JF cells were fixed and processed for fluorescent immuno-staining of endogenous CAMKV, again showing a widespread cytosolic localization in interphasic cells. **C**. mCherry-CAMKV JF and NGP cells were left untreated (left panels) or treated with the indicated final concentrations of Harmine for 24. Harmine treatments induced a relocalization of CAMKV from a homogeneous distribution into numerous aggregates (see Supplementary videos and fig. 7). **D**. Same cells from B. demonstrating a clear mitotic spindle localization of CAMKV in cells undergoing cell division. **E**. *Left Panel:* Same cells from B and D were processed for fluorescent immuno-staining of endogenous phospho-AKT1S1 (Thr 246), showing a clear mitotic spindle pole localization in cells undergoing cell division. Scale bar = 10 µm. *Right Panel:* Kaplan-Meier plot for the correlation between AKT1S1 expression and NB patient survival probability in the Cangelosi 786 cohort, as obtained from the ‘R2: Genomics Analysis and Visualization Platform (https://hgserver1.amc.nl/cgi-bin/r2/main.cgi).

## Discussion

In the present study we provide robust evidence demonstrating a role for the DYRK3 kinase as a critical modulator of Neuroblastoma cell proliferation and tumor growth. Our results suggest that this activity might be associated to the ability of DYRK3 to interact with and phosphorylate CAMKV, a protein abundantly expressed in high-risk NB and -like DYRK3-strongly correlated to a worse patient survival probability. Unfortunately, specific DYRK3 inhibitory compounds are currently lacking. Our data is consistent with a role for DYRK3 in the regulation of CAMKV partitioning into liquid-liquid phase separated biomolecular condensates, possibly due to the presence of an intrinsically disordered region in CAMKV’s C-terminal half, containing 7 tandem repeats of an octapeptide motif that may be directly phosphorylated by DYRK3, and thus, likely regulating the relocalization of CAMKV into specific membraneless organelles.

CAMKV remains a largely understudied protein, and this lack of knowledge is extensive to its localization and functions. Some authors have suggested membrane and/or vesicle localization, although the data supporting such features warrant further corroboration. Moreover, membrane localization could indeed result from cell type-, species- or context-specific characteristics. We provide evidence demonstrating that CAMKV is homogeneously distributed in the cytosol of interphasic NB cells. When NB cells enter the cell cycle, CAMKV levels increase and becomes relocalized to the mitotic spindle. Given the reported localization of DYRK3 to the mitotic spindle poles (Rai, 2018) and our observation of endogenous CAMKV and phospho-AKT1S1 (Thr246) in the mitotic spindles of NB cells, we hypothesize a critical role for this novel DYRK3/CAMKV/ AKT1S1 module in the assembly, maintenance or dissolution of the mitotic spindle of dividing NB cells. Finally, although CAMKV was originally predicted to lack a kinase activity, this feature has not been properly addressed. Given that CAMKV is almost exclusively expressed by NB cells and tumors, we speculate that novel small molecule inhibitors to specifically suppress the DYRK3/CAMKV module could constitute an innovative therapeutic strategy to fight high-risk neuroblastoma in a very precise and effective manner, a topic that will be the subject of our future efforts.

## Material and Methods

### Cell lines and culture

The neuroblastoma cell line SJNB-JF-G12 (JF) was originally established in 1979 from a patient with disseminated neuroblastoma and was a kind gift from Dr. Malcom Brenner. The NGP cell line was obtained from DSMZ; SH-SY5Y cells were obtained from ATCC. All cell lines were cultivated at 37 °C with 5% CO_2_. Cell lines were cultured according to vendors’ recommendations and passaged no more than 8 times. JF cells were grown in RPMI 1640 + 10% h.i. FBS + 4mM L-Glutamine + pen/strept. NGP cells were grown in DMEM (4.5 g/L glucose) + 10% h.i. FBS + 4mM L-Glutamine + pen/strept. SH-SY5Y cells were grown in DMEM/F12 (1:1) + 10% h.i. FBS + 4mM L-Glutamine + penicillin/streptomycin. All cell lines were tested for Mycoplasma every 2 months using e-Myco™ Mycoplasma PCR Detection Kit (Bulldog Bio #25235), according to the manufacturer’s instructions. All cell lines were authenticated by STR analysis (ATCC).

### Reagents

Harmine and GSK-626616 were obtained from MedChemExpress (HY-N0737A and HY-105309, respectively) and resuspended to a 10mM stock in DMSO for *in vitro*/cell culture-based assays. DAPI was from Millipore Sigma (#268298). Primary antibodies from Cell Signaling Technology were: mCherry (#43590); Δ-actin (#4970); vinculin (#13901), Phospho-PRAS40 (Thr246) (#2997) and HA-tag (#3724). Other primary antibodies were: anti-GAPDH from Millipore Sigma (MAB374), anti-DYRK3 from Aviva Systems Biology (ARP30648_P050), anti-CAMKV from Sino Biological (12243-T26). Mouse anti-rabbit (211-035-109) and goat anti-mouse (115-035-146) HRP-conjugated secondary antibodies were from Jackson ImmunoResearch Laboratories Inc. Purified human recombinant DYRK3 was obtained from Sino Biological (#10726-H20B). Dynabeads Protein A-magnetic beads for immunoprecipitation were obtained from Invitrogen (#10001D). Lipofectamine 2000 transfection reagent was from Invitrogen (#11668027).

### Western blot

Western blot analysis was conducted using standard methods (19). Briefly, cells grown to a 60-80% confluency were lysed in radioimmunoprecipitation assay (RIPA) lysis buffer (Prometheus Protein Biology Products #18-416) supplemented with Protease and Phosphatase Inhibitor Cocktails (Pr/Ph-ICs; Pierce, Thermo Scientific A32955 and A32957). Lysates were sonicated on ice, centrifuged at 15,000×g at 4 °C for 20 minutes and the soluble protein fraction was collected. Protein extracts were quantified using a Pierce BCA Protein Assay Kit (Thermo Scientific #23227). A total of 30-50 μg of proteins were separated via SDS-PAGE using Novex™ WedgeWell™ 4-20%, Tris-Glycine Mini Protein Gels (Invitrogen, Thermo Scientific XP04202BOX) and blotted onto a PVDF membrane using an iBlot transfer system and transfer stacks (Invitrogen, Thermo Scientific IB401001). Proteins were detected using SuperSignal™ West Pico PLUS Chemiluminescent Substrate (Thermo Scientific 34580). A ChemiDoc MP Imaging System (Bio-Rad) was used for chemiluminescent detection and analysis.

### Immunoprecipitation

For soluble cell lysates, cells were washed twice in PBS and lysed in IP buffer (50 mM Tris-HCl [pH 7.4], 150 mM NaCl, 2 mM EDTA, 1% NP-40, 10% glycerol, Pr/Ph-ICs). Lysates were clarified by centrifugation and protein quantification with the BCA assay kit (Pierce). One mg of total protein/sample were incubated for 6 h at 4°C with protein A/G-magnetic beads (Dynabeads, Invitrogen) prebound with 5 μg of anti-mCherry antibody, and then beads were washed with IP buffer. Finally, both lysates (input) and the immuno-precipitates were resuspended in 5X LB and analyzed by Western blotting.

### *In vitro* kinase assay

For *in vitro kinase* (IVK) from immunocomplexes, cell lysates were prepared in IP lysis buffer (50 mM HEPES [pH 7.4], 75 mM NaCl, 1 mM EDTA, 1% NP-40, Pr/Ph-ICs). Cell lysates were incubated overnight at 4°C with anti-mCherry bound to protein A-magnetic beads (Dynabeads, Invitrogen). Immunocomplexes were washed 4 times with IP and twice time with kinase buffer (25 mM Hepes pH 7.4, 5 mM MgCl2, 5 mM MnCl2, 0.5 mM DTT). Anti-mCherry immuno-complexes were split into 3 aliquots: 5% for Western blotting and 2 aliquots of 45% for IVK assay, with and without human recombinant DYRK3. Immunocomplexes were incubated for 20 min at 30°C in 30 μl of kinase buffer with a final concentration of 50 μM ATP and [γ-^32^P] ATP (1 × 10^−2^ μCi/pmol). Reactions were stopped by adding 5X LB, and samples were resolved by SDS-PAGE and then stained with Coomassie blue. ^32^P incorporation was detected by autoradiography of dried gels.

### Orthotopic xenografts-renal capsule injection

One million Tet-shDYRK3 JF cells suspended in 0.1 ml of PBS were surgically implanted in the left renal capsule of NSG immunodeficient mice (The Jackson Laboratory #005557). After days, mice were randomly divided into ‘control’ (untreated) group vs. ‘doxy’ group, receiving 1mg/ml doxycycline in the drinking water for 3 additional weeks. The body weight of mice was monitored weekly. At the end of the treatment, all mice were euthanized. Tumors and the right kidneys (control) were dissected and weighed. All procedures were approved by the Institutional Animal Care and Utilization Committee (IACUC) at UMass Chan Medical School and according to our IACUC approved protocol (A3306-01).

### RT-qPCR

Total RNA was extracted from cells using the Quick-RNA MiniPrep Kit (Zymo Research #11-327) following the manufacturer’s protocol. cDNA was synthesized from 0.5 ug/sample of total RNA with the ABScript II RT Mix (ABclonal RM21452) according to the manufacturer’s instruction. The cDNA was amplified in 96-well reaction plates with a Universal SYBR Green Fast qPCR Mix (ABclonal RM21203) on a QuantStudio 3 real-time PCR system (Applied Biosystems, Thermo Fisher Scientific). The sequences of forward and reverse primers are available upon request. The relative level of target transcripts was calculated from triplicate samples after normalization against human TBP and/or GAPDH transcripts. Dissociation curve analysis was performed after PCR amplification to confirm primers’ specificity. Relative mRNA expression was calculated using the ΔΔC_T_ method.

### Cell proliferation analysis

Cell proliferation of indicated conditions (untreated vs. shRNA-expressing cells) was measured as the relative whole-well confluency of 96-well culture plates using a Celigo Imaging Cytometer (Nexcelom Bioscience LLC). Briefly, ∼1000 cells/well were plated on day -1 and incubated for 24h to allow the cells to attach and recover. The following day each well was imaged and analyzed for relative confluence (day 0). After imaging, cells were treated with vehicle (DMSO, ‘untreated control’) or the indicated final concentrations of Harmine or Doxycycline for the indicated times. Relative confluence was subsequently analyzed every 24/48hs until untreated control wells reached 70-90% confluency. Each condition was done in 3-6 replicates per experiment. Medium +/- treatment was changed every 48-72h. Cell proliferation was plotted as the time-dependent change of the average relative confluency for each condition using GraphPad Prism 8 software.

### Immunofluorescence analysis

Standard immunofluorescence techniques were used as recommended in the Cell Signaling Technology protocols webpage. Briefly, cells were fixed in 4% paraformaldehyde in PBS for 15 min at room temperature or in 100% ice-cold methanol for 15 min at 4ºC (for anti-CAMKV), washed 2X in PBS (5 min/wash) and 2X with 0.2% Triton X-100 in PBS (‘PBS-T’). Cells were blocked in 2% BSA-PBS-T (blocking buffer) for 60 min at 4ºC and incubated with primary antibodies diluted in blocking buffer overnight at 4ºC. Then, 4X washes in PBS and 1h incubation with secondary antibody Alexa 488 goat anti-rabbit (Invitrogen) followed by 4X additional washes in PBS. Images were obtained in an Echo Revolve fluorescence microscope (Bico) or in a Zeiss LSM700 confocal microscope (UMass Chan Medical School). Images of mCherry-expressing constructs was obtained from *in vivo* microscopy with the Echo Revolve fluorescence microscope.

### Lentivirus preparation and infection

HEK-293T cells were maintained at 37ºC in Dulbecco’s modified Eagle medium (DMEM), supplemented with 10% FCS and antibiotics (100 units/ml penicillin and 100 μg/ml streptomycin). Cells were transfected with pVSV-G (Stewart *et al*., 2003) and pCMVΔR8.91 (Zufferey *et al*., 1997), together with the pLKO.1-puro non-targeting vector (Sigma Mission clone SHC016; ‘shControl’) or the pLKO.1-puro-shRNA vectors to target DYRK3 or CAMKV (Sigma Mission clone number available upon request, obtained from the UMass Chan Medical School RNAi core facility) using Lipofectamine™ 2000 reagent (Invitrogen) as recommended by the fabricant, and following the recommendations of The RNAi Consortium (TRC) laboratory protocols with slight modifications. Twelve hours after transfection the medium was replaced by DMEM, supplemented with 30% FCS and antibiotics which. Cell supernatants were harvested every 24 hs, replacing with fresh medium and stored at 4ºC until collection of the last harvest (at 72 hs). At this point, the consecutive harvests were pooled, filtered through 0.45 µm filters and split in 3-5 mL aliquots, which were stored at -80ºC. NB cells were infected with shControl or shRNA lentiviral particles by adding a 1:1 mix of medium:viral supernatant for 24-48 hs. Puromycin selection (2 μg/ml) was applied for 2-3 days and always compared to non-transduced control cells, which generally died within the first 24 hs. Target downregulation was confirmed by Western Blot and/or RT-qPCR. For mCherry overexpressing constructs the same lentiviral production strategy was followed, using a lentiviral expression construct instead of the pLKO.1-puro shRNA vectors. pLV-mCherry-CAMKV (WT) under the medium-strength promoter EFS (Human eukaryotic translation elongation factor 1 α1 short form) was custom-designed and ordered from VectorBuilder. mCherry-only or mCherry-CAMKV-ΔIDR variants were cloned from the original pLV-mCherry-CAMKV construct by standard molecular biology techniques. For the tetracycline-inducible shDYRK3 system, the Tet-pLKO-puro vector was obtained from Addgene (#21915) and cloning of the shDYRK3-2 hairpin (TRCN0000000647) was performed as recommended in the Tet-pLKO manual (available at www.addgene.org). Cells expressing mCherry were purified by FACS at the UMass Chan FACS core facility.

### Statistical Analyses

All quantitative data points represent the mean of three independent experiments performed in 3 or more replicates with standard deviation (S.D). Statistical analysis was performed using t-test or two-way ANOVA (GraphPad Software, Inc., La Jolla, CA).

