## Supplemental figures 1-9 for "A novel druggable DYRK3/CAMKV signaling module for Neuroblastoma tumor growth inhibition"

### Supplementary materials

Suppl. Fig. 1

| blue=worse | Brain Cohort (number of patients) | | | | | | | score |
|---|---|---|---|---|---|---|---|---|
| red=better | Madhavan 550 (REMBRANDT) | | | | TCGA 532 (Glioma) | French 284 (Glioma) | Kawaguchi 50 (Glioma) | |
| DYRK1A | 4.20E-04 | 4.28E-01 | 1.90E-02 | 8.39E-01 | 7.40E-02 | | 7.07E-01 | 11 |
| DYRK1B | | | | | | | | 0 |
| DYRK2 | 2.20E-04 | 3.30E-03 | 3.79E-01 | | | 5.70E-02 | | 11 |
| DYRK3 | 9.00E-13 | | | | 1.38E-08 | 8.12E-05 | 4.04E-03 | -44 |
| DYRK4 | | | | | | | | 0 |

Supplementary fig.1
Bonferroni adjusted p-values for the correlations between DYRK family members expression and Brain tumors patient survival probability. Values highlighted in blue denote significant negative correlations (higher expression = reduced survival probability ‘worse’), while values highlighted in red denote positive correlations (higher expression = increased survival probability ‘better’). Blue/red fonts denote non-significant values (bonf. p-val > 5.00E-02). Scores are arbitrary units based on the number of datasets showing one or more significant correlations for a given gene, and the value of such significance. We arbitrarily define that a gene must be significantly correlated to either a better or a worse outcome in at least 2 independent cohorts, and obtain a minimum of 20 points (either positive or negative, respectively), to be considered a significant hit.

Suppl. Fig. 2

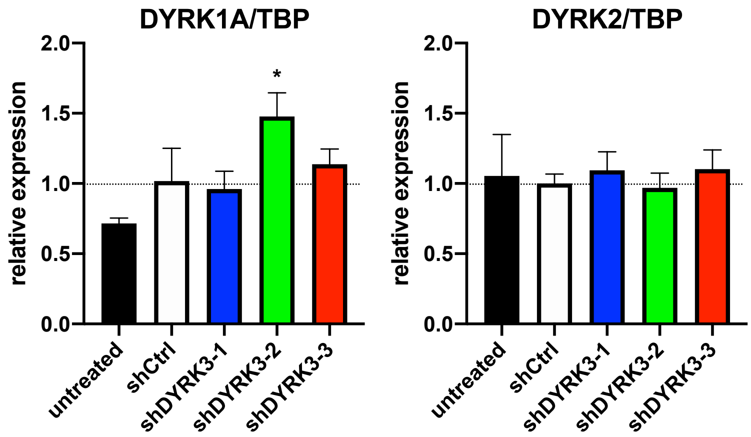


Supplementary fig. 2
The same cDNA samples from Figure 1B were subjected to qRT-PCR analysis of DYRK1A and DYRK2 expression (vs. the housekeeping gene TBP). No dramatic changes were observed in the expression of other family members upon downregulation of DYRK3.

Suppl. Fig. 3

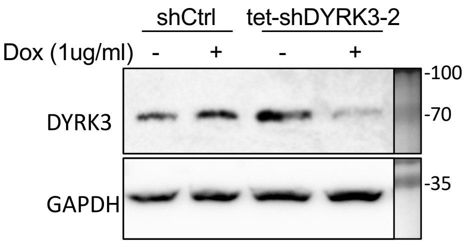

Supplementary fig. 3
The same cells from Figure 1C were processed in parallel for protein extraction and Tetracycline-inducible shRNA system validation by WB. DYRK3 protein expression was efficiently down-regulated upon doxycycline treatment of Tet-shDYRK3-expressing cells.

Suppl. Fig. 4

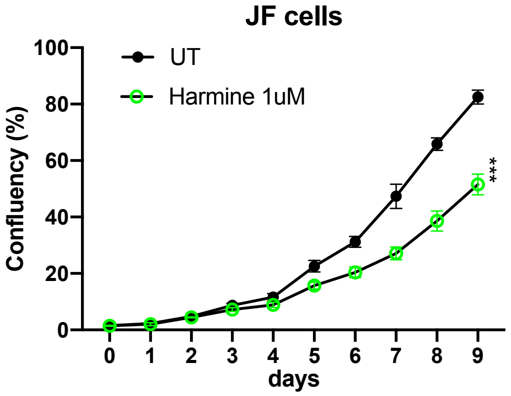


Supplementary fig. 4
Related to Figure 1D. JF cells were plated for proliferation assays in the absence (DMSO) or presence of 1 μM Harmine.

Suppl. Fig. 5

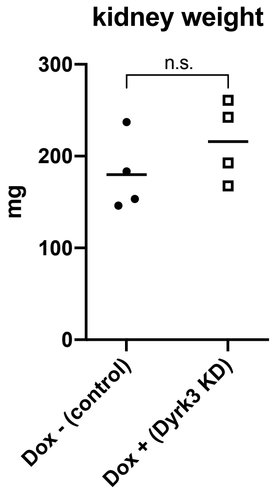


Supplementary fig. 5
Related to Figure 2. Contralateral healthy kidneys from *in vivo* Tet-shDYRK3 JF tumor formation experiment showed no significant differences in size/weight.

Suppl. Fig. 6

PONDR: Predictor Of Natural Disordered Regions

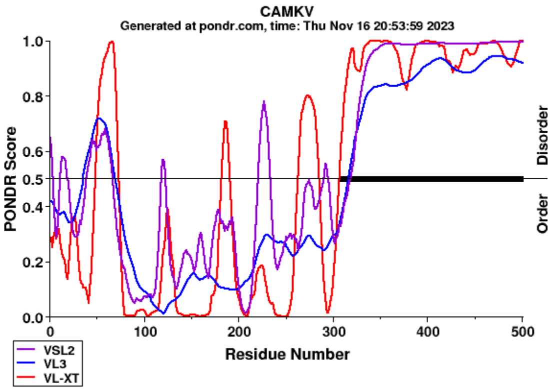


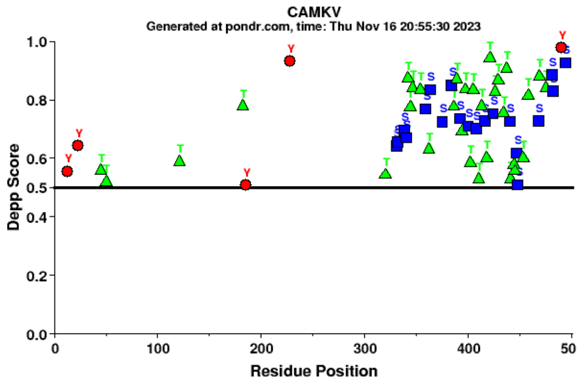


PrDos: Protein DisOrder prediction System

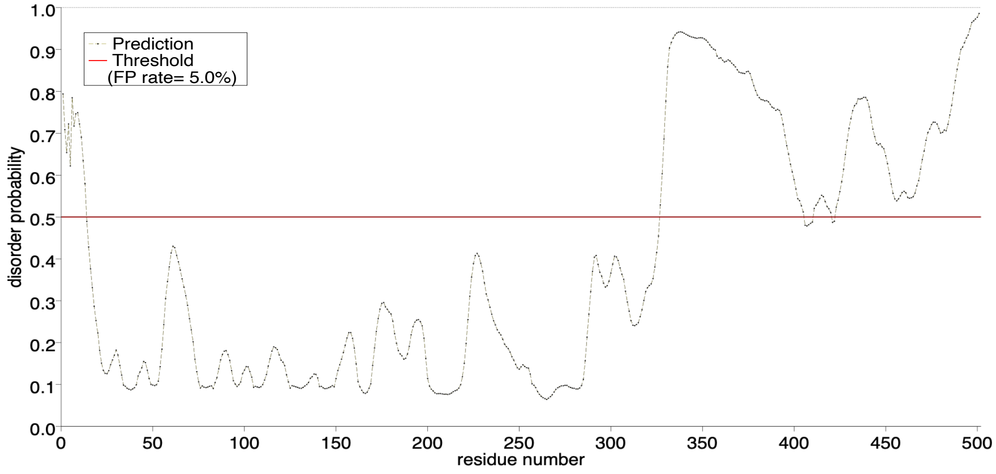


Supplementary fig. 6

Related to Figure 4A. Human CAMKV protein sequence was analyzed with the online PONDR (Predictor Of Natural Disorered Regions; www.pondr.com) tool, suggesting a highly disordered C-terminal region of ~200 amino-acids (upper left panel). We also used the Disordered Enhanced Phosphorylation Predictor (DEPP) tool from the PONDR webpage. DEPP uses disorder information to improve the discrimination between phosphorylated and non-phosphorylated sites.

Suppl. Fig. 7

NGP mCherry-CAMKV

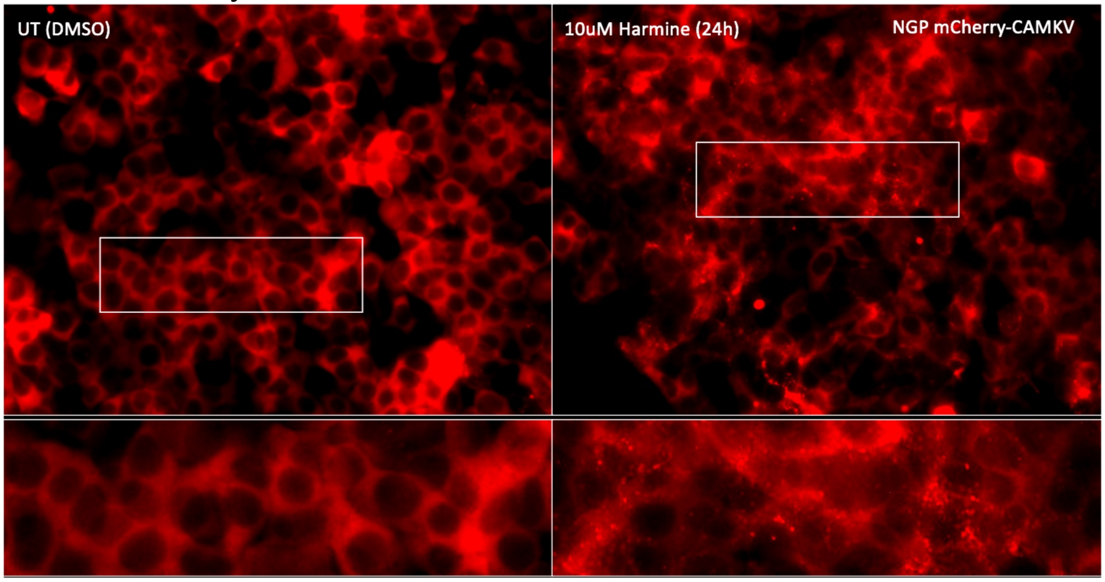


Supplementary fig. 7

Related to Figure 5C. mCherry-CAMKV NGP cells were left untreated (DMSO; left panels) or treated with the indicated final concentrations of Harmine for 24. Harmine treatments induced a relocalization of CAMKV from a homogeneous distribution into numerous aggregates.

Suppl. Fig. 8

NGP mCherry-CAMKV

Untreated (DMSO)

10 μM GSK-626616

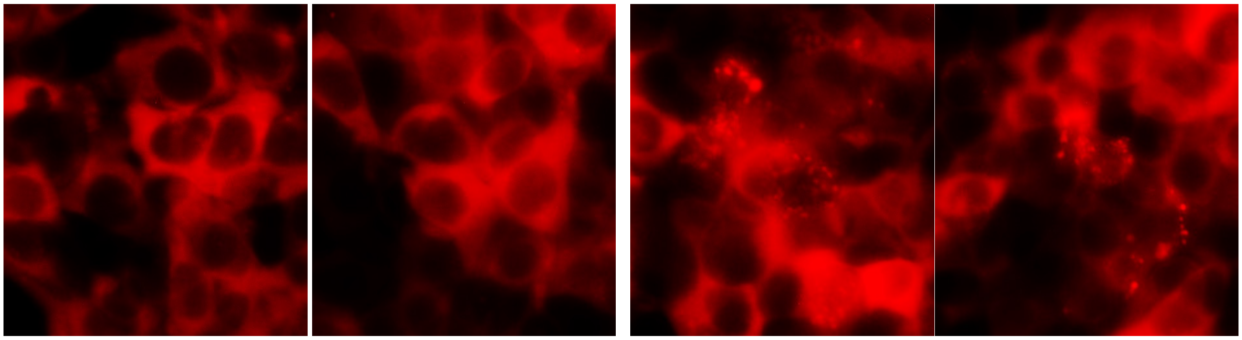

Supplementary fig. 8

Related to Figure 5C. mCherry-CAMKV NGP cells were left untreated (DMSO; left panels) or treated with the indicated final concentrations of the DYRK3 inhibitor GSK-626616 for 24. GSK-626616 treatment induced a relocalization of CAMKV from a homogeneous distribution into aggregates.

Suppl. Fig. 9

JF anti-CAMKV IF (endogenous) - confocal

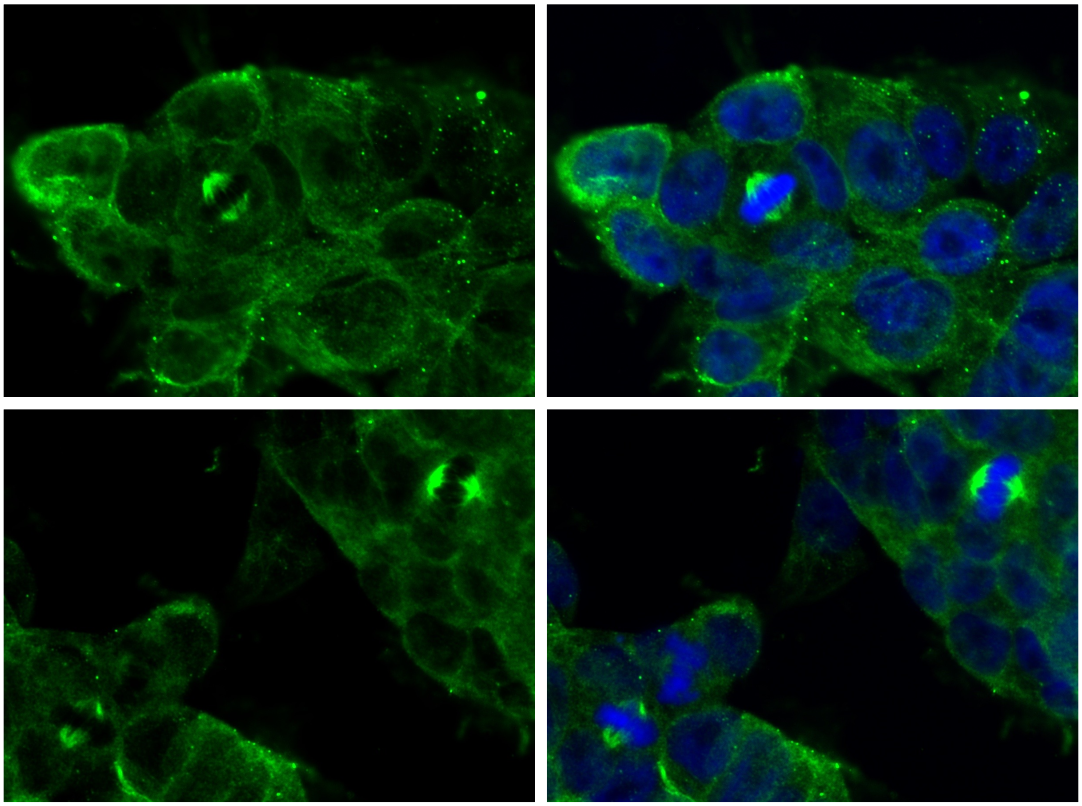

Supplementary fig. 9

Related to Figure 5D. JF cells were fixed and processed for fluorescent immuno-staining of endogenous CAMKV by laser confocal microscopy, confirming a clear mitotic spindle localization of CAMKV in cells undergoing cell division.
